# Immune Remodeling and Dysbiosis May Distinguish the Microenvironments of Gastric Adenocarcinoma and Peritumoral Tissue

**DOI:** 10.1101/2025.08.27.672687

**Authors:** Ronald Matheus da Silva Mourão, Juliana Barreto Albuquerque Pinto, Jéssica Manoelli Costa da Silva, Daniel de Souza Avelar da Costa, Valéria Cristiane Santos da Silva, Ana Karyssa Mendes Anaissi, Samia Demachki, Williams Fernandes Barra, Fabiano Cordeiro Moreira, Paulo Pimentel de Assumpção

**Affiliations:** Núcleo de Pesquisas em Oncologia, Federal University of Pará, Belém, PA, Brazil; Graduate Program in Genetics and Molecular Biology, Federal University of Pará, Belém, PA, Brazil

**Keywords:** Gastric cancer, Tumor microenvironment, Microbiome, Microbiome-immune interactions

## Abstract

The gastric tumor microenvironment is dynamically shaped by the interactions between the local microbiota and the host immune system, although the functional integration of these elements remains incompletely understood. In this study, we characterized microbial diversity, immune cell composition, and immune-related gene expression profiles in samples of gastric adenocarcinoma (GAC) and adjacent peritumoral tissue (PTT), aiming to elucidate their functional organization. A total of 106 samples of 75 patients were analyzed using bulk RNA-Seq expression profiling, immune deconvolution, and bacterial taxonomic reconstruction. While alpha diversity remained preserved between GAC and PTT, distinct compositional differences emerged: GAC was enriched with *Pseudomonadota, Enterobacteriaceae*, and *Escherichia*, whereas PTT exhibited a predominance of *Helicobacteraceae* and *Helicobacter*. Immune deconvolution revealed an expansion of cancer-associated fibroblasts (CAFs) and mast cells in GAC, correlated with higher expression levels of *TGFB1* and *FOXP3*, while neutrophils and B cells predominated in PTT. Integrated analysis demonstrated that GAC formed dense and cohesive networks connecting pro-inflammatory bacteria, activated immune cells, and inflammatory genes such as *IL1B, CXCL8*, and *IFNG*. In contrast, PTT exhibited dispersed networks and negative correlations, suggesting a less structured, tolerogenic environment. Our findings indicate that gastric cancer progression involves not only compositional shifts in microbiota and immune cells but also the active construction of functionally integrated inflammatory networks, providing new insights into potential therapeutic targets at the microbiome-immune interface.

## 1 Introduction

Gastric adenocarcinoma (GAC) is a multifactorial epithelial malignancy whose progression involves not only intrinsic genetic alterations within tumor cells but also progressive reprogramming of the surrounding tissue microenvironment[1–3]. Within this context, the immune-inflammatory axis and the influence of the microbiome have emerged as central elements in the transition from inflamed mucosa to established tumor states[4, 5]. Chronic activation of the immune system, phenotypic remodeling of fibroblasts, and the presence of specialized bacterial consortia collectively contribute to the creation of a permissive environment for carcinogenesis and immune evasion[6].

Functional compartmentalization of the gastric microenvironment - segregating inflammatory responses, adaptive immunity, and microbial stimuli - is a critical feature of tissue homeostasis[7]. As tumor progression advances, this compartmentalized architecture tends to collapse, fostering the overlap of chronic inflammation, immunosuppression, and bacterial dysbiosis[2]. Previous studies have demonstrated that immune infiltration in GAC is marked by signs of functional exhaustion and a predominance of tolerogenic inflammatory profiles, in contrast to the more balanced environment observed in peritumoral tissues[8]. However, the spatial and functional dynamics of microbiome, immunity, and gene expression interactions during gastric tumor progression remain poorly understood.

The gastric microbiome, traditionally associated with *Helicobacter pylori*, is now recognized as a broader and more dynamic ecosystem capable of modulating inflammatory pathways, altering local cellular profiles, and influencing tumor evolution[9–11]. Certain microbial communities promote immunosuppressive environments, whereas others drive pro-inflammatory activation, directly reshaping the functional architecture of the microenvironment[12]. The tripartite interaction between epithelium, immunity, and microbiota thus emerges as a key axis in configuring the functional heterogeneity of gastric tissues.

Understanding how microbial and immune networks organize - or become disorganized - during GAC progression is critical for identifying therapeutic intervention points[13]. This study aims to delineate the functional integration among the microbiome, cellular composition, and gene expression profiles in gastric adenocarcinoma and adjacent peritumoral tissue, characterizing the structural transitions of the microenvironment associated with tumor progression.

## 2 Material and Methods

### 2.1 Sample Characterization and Ethical Considerations

In this study, tumor and adjacent peritumoral tissue (PTT) samples were collected from patients diagnosed with GAC, the most common type of gastric cancer. A total of 75 patients were analyzed, comprising 62 GACs tissues and 44 PTT samples. The cohort included 29 female and 45 male patients and 1 patient with unreported gender. GAC samples were staged according to the ypTNM classification: Of the patients for whom staging information was available, 6 were classified as stage I, 18 as stage II, 32 as stage III, and 3 as stage IV. Recruitment and sample collection were conducted between July 2, 2022, and July 6, 2023, at the João de Barros Barreto University Hospital in Belém, Brazil. The study objectives were clearly explained to all participants, who provided written informed consent. The study was conducted in accordance with the Declaration of Helsinki and approved by the Ethics Committee of the João de Barros Barreto University Hospital (approval number: 47580121.9.0000.5634).

### 2.2 RNA Extraction and Quality Assessment

Approximately 50-100 mg of tissue from each sample were macerated, followed by the addition of 1 mL of TRIZOL® reagent to facilitate RNA extraction. The integrity and concentration of total RNA were evaluated using Qubit 4.0 (Thermo Fisher Scientific) and NanoDrop ND-1000 (Thermo Fisher Scientific) fluorometers. Optimal criteria for total RNA integrity were considered met when samples exhibited an A260/A280 ratio between 1.8 and 2.2, an A260/A230 ratio greater than 1.8, and an RNA Integrity Number (RIN) ≥ 5. This threshold was selected to accommodate the inherent variability in RNA quality from clinical tissue samples, ensuring the inclusion of a representative cohort while maintaining data reliability.

### 2.3 cDNA Library Construction and Sequencing

The TruSeq Stranded Total RNA Library Prep Kit with Ribo-Zero Gold (Illumina) was used to remove cytoplasmic and mitochondrial rRNA. Libraries were processed using the NextSeq® 500 High Output V2 kit - 150 cycles (Illumina), following the manufacturer’s specifications. After library construction, a new assessment of RNA integrity was performed using the 2200 TapeStation System (Agilent). The cDNA libraries were then loaded onto the Illumina NextSeq sequencing platform and sequenced in paired-end mode.

### 2.4 Quality, Alignment, Quantification and Transcriptome Expression

Read quality was assessed using FastQC (v0.11.9), and low-quality reads and adapter sequences were removed with Trimommatic, applying a minimum Phred quality threshold of QV15. QV15 was selected as a pragmatic threshold, given that Salmon’s k-mer-based pseudoalignment is robust to moderate base quality variation. Filtered reads were quantified at the transcript level using Salmon (v1.5.2)[14] against the human transcriptome reference (hg38). Transcript abundances were imported using the Tximport[3], and a DESeq2[15] object was created to normalize gene expression levels, accounting for tissue type (GAC or PTT) and sequencing batch effects. Variance-stabilized (VST) and batch-corrected expression values were used for subsequent analyses.

### 2.5 Selection of Immune-Related Genes

A curated panel of immune-related genes was assembled to investigate key processes within the tumor microenvironment and host-microbiome interactions. The selection was informed by comprehensive immunological databases, such as MSigDB, and was further refined based on our group’s previous unpublished study. The panel included classical immune checkpoints (*PDCD1, CD274, CTLA4, LAG3, HAVCR2, CD47*), pro-inflammatory cytokines and mediators (*IFNG, TNF, IL6, IL1B, CXCL8, CCL2, CCL5*), regulatory and immunosuppressive markers (*IL10, TGFB1, FOXP3*), macrophage and myeloid cell markers (*CD163, CD68, CD86, CD83*), signaling molecules involved in immune activation and regulation (*STAT3, MYD88, NFKB1*), as well as genes associated with antigen presentation (*B2M, HLA*.*A*), epithelial plasticity (*SOX9*), angiogenesis (*VEGFA*), and mucosal immune defense (*PIGR*). This focused selection allowed a comprehensive assessment of inflammatory activation, immune regulation, stromal remodeling, and adaptive responses.

### 2.6 Microbiome

Microbiome characterization was performed by taxonomic classification of RNA-Seq reads using Kraken2 (v2.1.4) [16] against the comprehensive PlusPF database. The primary focus of this study was to quantify bacterial microbiome expressions associated with GAC. To achieve this, relevant bacterial genomes were obtained from the RefSeq database, and Salmon (v1.10.1) was used to quantify expression by aligning reads against these genomes. The resulting expression counts were used to estimate bacterial abundance.

To prioritize the most representative bacterial genera across samples, we computed an abundance score that integrates both dominance and prevalence (equation 1) [17]. For each sample, genera were ranked in descending order based on their absolute abundance. The mean rank of each genus across all samples in which it was detected 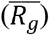 was then multiplied by a frequency-based penalty factor (1.1 - *f_g_*), where *f_g_* represents the proportion of samples in which that genus was present. The constant 1.1 was introduced to avoid disproportionately penalizing highly prevalent genera, ensuring that those consistently detected and highly ranked retained meaningful scores.

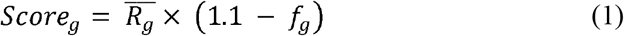

Based on this criterion, the 15 most abundant genera were selected to effectively represent the most relevant taxa for downstream analyses. Additionally, genera with recognized roles in microbiome-immune interactions and tumor biology - such as *Parvimonas, Peptostreptococcus, Campylobacter, Actinomyces, Escherichia, Klebsiella, Streptococcus, Helicobacter, Prevotella, Halomonas, Pseudomonas, Sphingomonas, Lactobacillus, Shewanella, Acinetobacter, Corynebacterium, Bacillus, Neisseria, Leptotrichia, Veillonella, Bacteroides, Faecalibacterium, Bifidobacterium, Chryseobacterium, Oscillospira, Haemophilus, Actinobacillus, Staphylococcus, Lactococcus, Porphyromonas, Propionibacterium*, and *Fusobacterium* - were also included, based both on published evidence and prior findings from our research group. Alpha diversity analysis was conducted using the Shannon, Chao1, and Observed indices, with group comparisons performed using the Wilcoxon rank-sum test and p ≤ 0.05. The *microbiome* package was used to estimate sample diversity between GAC and PTT groups.

### 2.7 Cellular Deconvolution

Cellular deconvolution was performed using three computational tools: CIBERSORT[18], quanTIseq[19], and EPIC[20]. The LM22 immune cell signature file was loaded, and gene expression data were normalized to generate a TPM (Transcripts Per Million) matrix. The CIBERSORT function was employed to estimate cell composition, and the run_quantiseq function was applied for additional cellular composition estimation in tumor samples. The EPIC package was used to calculate cellular fractions across the samples. The results from each tool were integrated to generate a comprehensive deconvolution table of immune cell fractions. Differences in cell proportions between GAC and PTT groups were assessed using the Wilcoxon test, with Benjamini-Hochberg correction for multiple testing (FDR ≤ 0.05).

### 2.8 Hierarchical Clustering

The dataset - comprising immune-related genes, inferred immune cell fractions, and relative abundances of bacterial genera - was transposed so that variables became rows, enabling the analysis of their similarities. To enable meaningful comparisons across variables with different scales and units, the data was first log2-transformed and subsequently standardized using z-scores.

The distance matrix between variables was calculated using standard Euclidean distance, and hierarchical clustering was performed using the Ward.D2 method, which minimizes the total variance within clusters. Cluster structure visualization was performed with the *fviz_dend* function from the *factoextra* package, using the “rectangle” type combined with the “layout.gem” radial layout.

Interpretation of the dendrogram focused exclusively on tree topology, considering the visual proximity of elements as indicative of relative functional similarity, as reflected in the original distance matrix. Closely clustered groups were interpreted as functionally related modules, whereas distant branches suggested differentiation among cellular, genetic, or microbial profiles within the GAC and PTT microenvironments.

### 2.9 Statistical Analyses

Correlations among bacterial abundance, gene expression, and immune cell fractions were evaluated using Spearman’s correlation. A threshold of |rho| > 0.3 and p ≤ 0.05 was retained in line with exploratory objectives and biomedical literature precedent. Additional statistical tests, such as the Wilcoxon test, were conducted to compare differences between experimental groups. Result visualizations, including boxplots, bar graphs, and significant correlations, were generated using the *ggplot2, ggcorrplot*, and *cowplot* packages.

## 3 Results

### 3.1 Microbiome

Alpha diversity analysis revealed no significant differences in Shannon indices between GAC and PTT (p = 0.6; Figure 1A), suggesting no notable variation in ecological heterogeneity between the two tissue types. Similarly, species richness measures did not differ between GAC and PTT (Figure 1B), supporting the notion of a global stability in microbial complexity.

**Figure 1.**
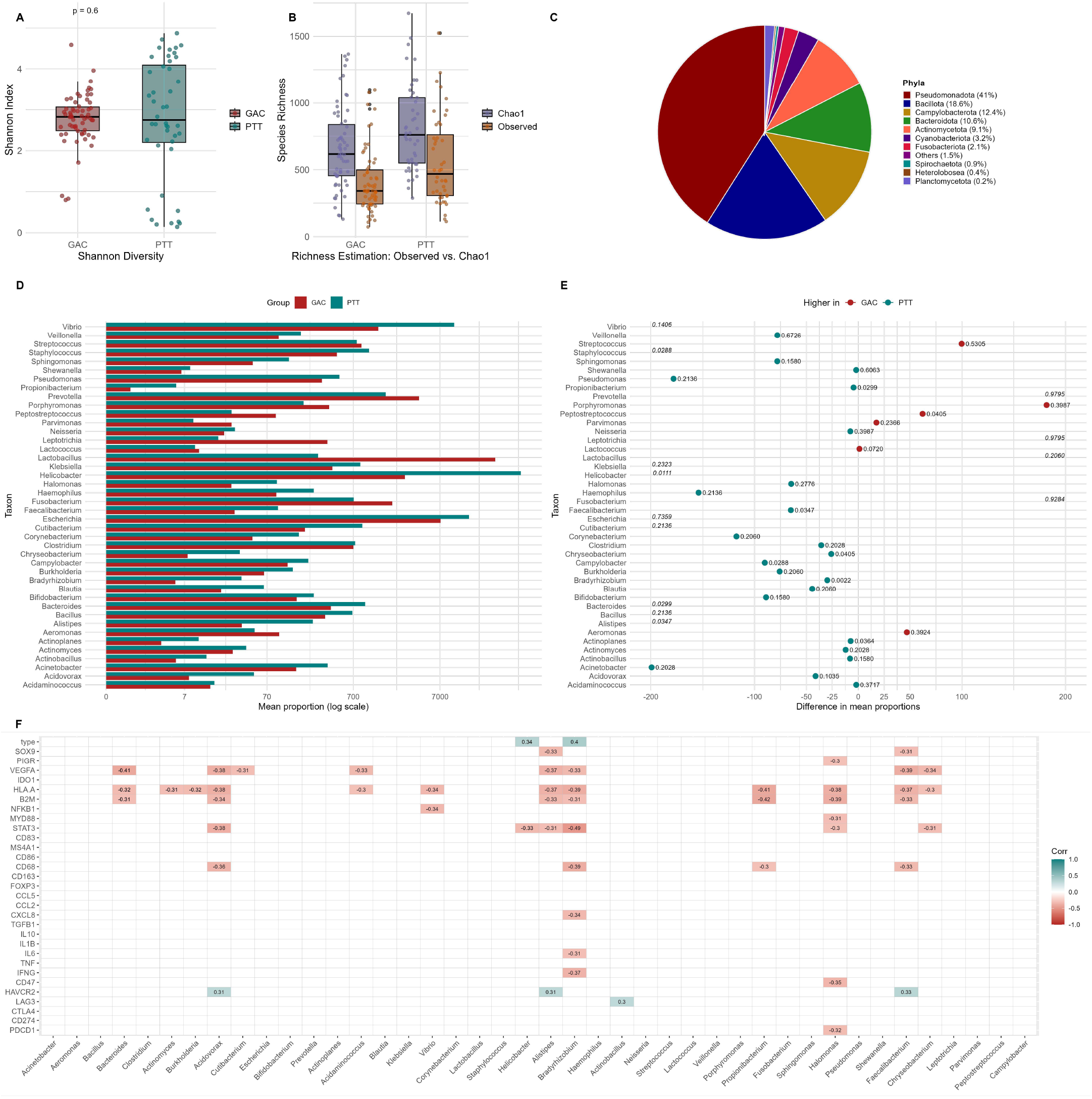
Bacterial microbiome analyses: (A) Shannon diversity index; (B) Species richness estimates (Observed and Chao1); (C) Relative abundance of major bacterial phyla; (D) Relative abundance of predominant bacterial genera; (E) Mean difference in genus abundance between GAC and PTT samples. (F) Correlation between bacterial abundance and the expression of immune response genes across all samples (GAC and PTT combined).

Regarding taxonomic composition, the phylum *Pseudomonadota* was the most dominant, accounting for 41% of the total, followed by *Bacillota* (18.6%), *Campylobacterota* (12.4%), and *Bacteroidota* (10.6%) (Figure 1C). At the family level, *Enterobacteriaceae* was the most abundant, representing 17.8% of the total microbiota and associated with GAC (66.4% of its fraction). In contrast, *Helicobacteraceae* was markedly more prevalent in PTT (76.3% of its relative abundance). *Lactobacillaceae* (4.4%) and *Streptococcaceae* (2.0%) were also more prominent in GAC (88.8% and 80.5%, respectively) (Figure 4A, see appendix I).

Among genera, *Escherichia* emerged as the most abundant (12.7% of the total), with 68% of its representation in GAC. Conversely, *Helicobacter* concentrated 76.3% of its abundance in PTT. Genera such as *Prevotella* (68.2% GAC) and *Lactobacillus* (88.7% GAC) were also more associated with the tumor environment, whereas *Cutibacterium* (39.4% GAC, 60.6% PTT) and *Rhizobium* (44.7% GAC, 55.3% PTT) displayed a more balanced distribution (Figure 1D).

Differential abundance analysis between genera highlighted several relevant disparities (Figure 1E). *Helicobacter* showed a highly significant difference (adjusted p < 0.001), confirming its greater prevalence in PTT. Other genera, including *Staphylococcus, Propionibacterium, Faecalibacterium, Chryseobacterium, Campylobacter, Bradyrhizobium, Bacteroides, Alistipes*, and *Actinoplanes*, were more abundant in GAC (adjusted p < 0.05). These differences remained significant after multiple-testing correction.

Correlation analysis between bacterial abundance and gene expression across all samples (GAC and PTT combined) revealed patterns characterized by negative associations (Figure 1F). Several genera exhibited inverse correlations with key genes involved in inflammatory responses and antigen presentation, including *Bacteroides* with *B2M, HLA*.*A*, and *VEGFA*; *Vibrio* with *NFKB1* and *HLA*.*A*; and *Bradyrhizobium* with *IFNG, IL6, CXCL8*, and *STAT3*. Some genera, such as *Acidovorax, Alistipes*, and *Faecalibacterium*, demonstrated mixed patterns, positively correlating with the immunoregulatory marker *HAVCR2* while negatively correlating with pro-inflammatory and angiogenic genes.

### 3.2 Immune Microenvironment Estimation

Cellular deconvolution revealed substantial quantitative differences between GAC and PTT (Figure 2A; Figure 5, see appendix). In GAC samples, epic_CAFs (p = 6.83 × 10^−9^), ep-ic_Macrophages (p = 1.53 × 10^−3^), quantiseq_Macrophages.M1 (p = 6.40 × 10 □ □), cibersort_Dendritic cells resting (p = 4.92 × 10^−2^), cibersort_Mast cells resting (p = 1.45 × 10^−2^), and epic_NKcells (p = 8.09 × 10^−3^) were significantly more abundant, delineating a tumor microenvironment enriched in stromal, myeloid, mast cell, and NK cell populations. In PTT samples, the most abundant populations were epic_Bcells (p = 1.63 × 10^−2^), quantiseq_B.cells (p = 1.63 × 10^−2^), epic_CD8_Tcells (p = 4.92 × 10^−2^), and cibersort_Neutrophils (p = 1.63 × 10^−2^), composing a more effector and inflammatory immune profile in this tissue.

**Figure 2.**
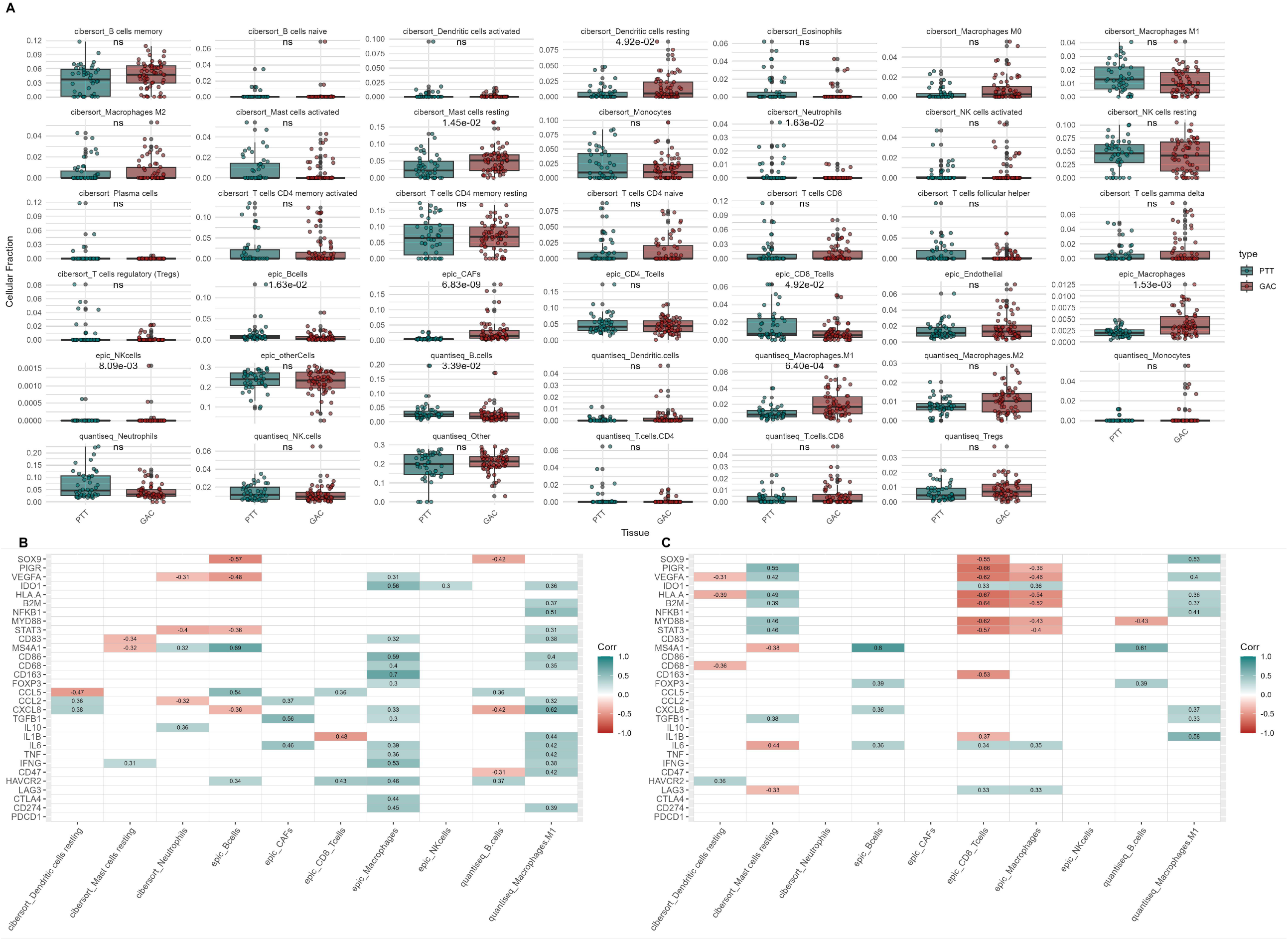
Immune deconvolution analyses GAC PTT: (A) Mean differences in immune cell proportions between GAC and PTT groups; (B) Correlations between immune cell proportions and immune response gene expression in GAC samples; (C) Correlations between immune cell proportions and immune response gene expression in PTT samples.

Global analysis of cellular proportions, considering all samples together, showed that cibersort_Mast cells resting comprised the largest fraction (30%), followed by quantiseq_B.cells (19.5%), epic_CAFs (13%), quantiseq_Macrophages.M1 (11.4%), and epic_CD8_Tcells (8.1%) (Figure 5, see appendix). These results indicate that in GAC and its PTT, the cellular landscape is dominated by stromal components, B lymphocytes, and myeloid cells.

Comparative analysis between GAC and PTT revealed notable structural contrasts. In GAC, a predominance of epic_CAFs (88.4%), cibersort_Dendritic cells resting (77.5%), epic_NKcells (78.1%), cibersort_Mast cells resting (68.7%), epic_Macrophages (72.8%), and quantiseq_Macrophages.M1 (75.8%) was observed, configuring a microenvironment dominated by stromal, myeloid, and mast cell populations. Although the overall proportion of epic_CD8_Tcells was lower in GAC compared to PTT, some tumor samples exhibited significant infiltration of cytotoxic T lymphocytes, suggesting intratumoral heterogeneity. In PTT, the most representative populations included cibersort_Neutrophils (83.2%), epic_Bcells (56.1%), and epic_CD8_Tcells (59.1%) (Figure 5, see appendix), reflecting a more effector-dominant microenvironment, characterized by greater infiltration of adaptive immune cells and localized inflammatory responses.

Correlation analysis between cellular composition and gene expression corroborated the structural patterns observed (Figures 2B-2C). In GAC samples, epic_CAFs showed positive correlations with immunosuppressive and extracellular matrix-modulating genes, notably *TGFB1* (ρ = 0.56) and *IL6* (ρ = 0.46). Quantiseq_Macrophages.M1 were strongly associated with inflammatory genes, including *CXCL8* (ρ = 0.62), *IL1B* (ρ = 0.44), *TNF* (ρ = 0.42), and *NFKB1* (ρ = 0.51), as well as with antigen-regulatory genes such as *CD86* and *CD83*. Epic_Bcells positively correlated with adaptive immune genes, particularly *MS4A1* (ρ = 0.69) and *PIGR* (ρ = 0.48), while also displaying negative correlations with inflammatory markers such as *CXCL8* (ρ = −0.36) and *VEGFA* (ρ = −0.48). Cibersort_Mast cells resting exhibited mixed patterns, positively associating with *IFNG* (ρ = 0.31) and negatively with *MS4A1* (ρ = −0.32).

In PTT samples, epic_Bcells maintained a strong positive correlation with *MS4A1* (ρ = 0.80) and correlated positively with inflammatory genes such as *IL6* (ρ = 0.36) and *CXCL8* (ρ = 0.36). Quantiseq_Macrophages.M1 showed positive associations with pro-inflammatory genes, notably *IL1B* (ρ = 0.58) and *CXCL8* (ρ = 0.37), as well as with regulatory genes such as *TGFB1* (ρ = 0.33) and *NFKB1* (ρ = 0.41). Cibersort_Dendritic cells resting demonstrated a positive correlation with *HAVCR2* (ρ = 0.36), while cibersort_Mast cells resting positively correlated with *TGFB1* (ρ = 0.38), *STAT3* (ρ = 0.46), and *VEGFA* (ρ = 0.42), and negatively with *IL6* (ρ = −0.44) and *CD274* (ρ = −0.35).

Overall, quantiseq_Macrophages.M1 and epic_Bcells were the cellular subsets that exhibited the highest number and intensity of correlations in both GAC and PTT. In GAC, associations were positive and related to inflammatory axes. In PTT, a combined pattern of positive correlations with both inflammatory (*IL1B, CXCL8*) and immunoregulatory (*TGFB1, HAVCR2*) genes were observed, reflecting a functionally more heterogeneous environment. Negative correlations, particularly involving cibersort_Dendritic cells resting and cibersort_Mast cells resting, were more frequent in PTT, suggesting localized patterns of immune modulation.

### 3.3 Integration of the Microbial, Immune and Genetic axis

Integrated analysis of the gene expression, immune cells, and microbiota in GAC and PTT revealed highly organized patterns of interaction, supported by robust correlations. In GAC, the formation of an immunoregulatory cluster composed of *IDO1, FOXP3, HAVCR2, IL10*, and *LAG3* stood out, reflecting the activation of immune suppression programs within the tumor microenvironment (Figure 3A). The correlation between *IDO1* and *FOXP3* (r = 0.57) and the coexpression of *PIGR* and *MS4A1* (r = 0.35) (Figure 3B) further reinforce the robustness of this regulatory signature. Methodological convergence in the detection of B cells, as evidenced by the strong correlation between *quantiseq_B*.*cells* and *epic_Bcells* (r = 0.74), adds additional consistency to these observations. In parallel, an inflammatory cluster consolidated the activation of effector immune response pathways, with associations between *IFNG* and *CD274* (r = 0.50), *CD86* and *CTLA4* (r = 0.63), and *CXCL8* and *IL1B* (r = 0.80), outlining an acute inflammatory environment associated with immune checkpoint activation.

**Figure 3.**
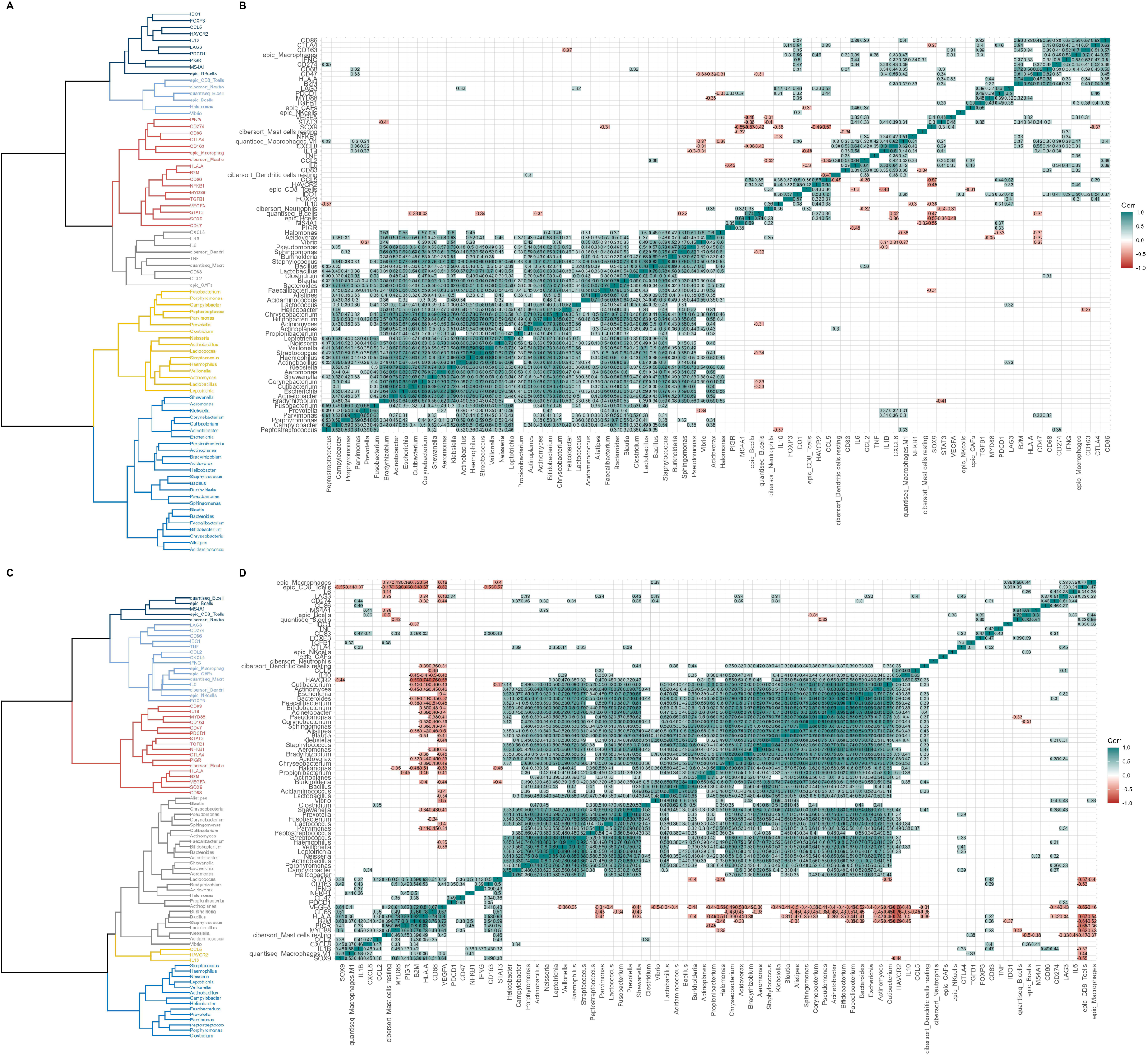
Integrated analysis of microbiome, immune cells, and gene expression in GAC and PTT: (A) Hierarchical clustering of bacterial genera, immune cell fractions, and immune-related genes in GAC samples; (B) Correlation matrix illustrating significant associations among bacterial abundance, immune cell fractions, and gene expression in GAC; (C) Hierarchical clustering of bacterial genera, immune cell fractions, and immune-related genes in PTT samples; (D) Correlation matrix illustrating significant associations among bacterial abundance, immune cell fractions, and gene expression in PTT.

Microbiome structuring revealed two distinct bacterial axes. A cluster of pathogenic oral bacteria, including *Fusobacterium, Prevotella, Porphyromonas, Haemophilus*, and *Veillonella*, exhibited strong co-occurrences, suggesting the formation of biofilms associated with tumor progression. Another cluster, composed of commensal and environmental bacteria such as *Blautia, Faecalibacterium, Bacteroides, Pseudomonas*, and *Escherichia*, indicated the coexistence of diverse ecological communities within the tumor microenvironment. *Helicobacter* showed relevant integration into both bacterial networks, linking to oral species (*Veillonella, Streptococcus*) and environmental species (*Pseudomonas, Escherichia*), suggesting its participation in complex microbial consortia within GAC.

In PTT analysis, the preservation of functional clusters was evident (Figure 3C). The potential adaptive immune cluster involving *quantiseq_B*.*cells, epic_Bcells*, and *MS4A1* stood out, indicating the persistence of adaptive responses in adjacent tissue, supported by strong correlations (r = 0.80 and r = 0.72) (Figure 3D).

Regarding the PTT microbiome, clusters of oral and environmental bacteria were evidenced by strong co-occurrences between *Alistipes* and *Faecalibacterium* (r = 0.87), *Fusobacterium* and *Leptotrichia* (r = 0.83), and *Prevotella* and *Neisseria* (r = 0.74). The integration of *Helicobacter* into PTT bacterial networks further supports the hypothesis of complex microbial adaptations occurring not only within tumors but also in adjacent tissues.

Although topological analysis revealed clustering in both tissue types, statistical validation indicated that not all observed proximities corresponded to robust associations, particularly in PTT; therefore, only functionally and statistically supported cores were emphasized in the results.

## 4 Discussion

This study revealed that although global bacterial microbiome diversity and richness are preserved between GAC and PTT, taxonomic composition and functional organization of the microenvironments diverge substantially[21]. These findings suggest that gastric tumor progression may involve not only a preservation of microbial heterogeneity, but also compositional shifts in bacterial consortia potentially driven by tumor-associated environmental pressures.

The preservation of Shannon, Observed, and Chao1 indices between GAC and PTT indicates that global ecological complexity is maintained during tumor progression. However, the redistribution of relative abundances - with enrichment of *Pseudomonadota* and *Bacillota* in GAC and Campylobacterota and Bacteroidota in PTT - points to selective ecological reprogramming. Conditions such as hypoxia, acidification, and nutritional imbalances in GAC likely function as selective pressures, favoring bacterial phyla more adapted to inflammatory and metabolically hostile environments[22].

At a finer taxonomic resolution, the greater abundance of *Enterobacteriaceae* and *Escherichia* in GAC, along with positive correlations with inflammatory genes such as *IFNG* and *CD86*, suggests that these bacteria may contribute to maintaining a chronic inflammatory state permissive to tumorigenesis[23]. In contrast, the predominance of *Helicobacteraceae* and *Helicobacter* in PTT, along with negative correlations with *IFNG* and *IL6*, may reflect a more regulatory environment, potentially characteristic of an early stage of immune escape[24]. We propose that the replacement of *Helicobacter* by pro-inflammatory bacterial consortia represents a critical transition in the remodeling of the gastric microenvironment.

Immune composition analysis further reinforced this interpretation. In GAC, an enrichment of epic_CAFs and cibersort_Mast cells resting was observed, whereas in PTT, quantiseq_B.cells, epic_Bcells, and epic_CD8_Tcells predominated[25–27]. The strong correlation between ep-ic_CAFs and *TGFB1* and *FOXP3* suggests that stromal fibrosis is integrated into immune suppression circuits within the tumor[28]. Conversely, the presence of adaptive B cells in PTT was corroborated by the strong correlation between quantiseq_B.cells and epic_Bcells (r = 0.74), indicating methodological consistency in the detection of this population. Importantly, this correlation reflects the identification of the same B cell population by distinct deconvolution methods (quanTIseq and EPIC)

Functional integration of the microbiome, cellular composition, and gene expression revealed the formation of highly organized pro-inflammatory axes in GAC[29]. The topological proximity of *Fusobacterium, Escherichia*, activated macrophages (quantiseq_Macrophages.M1), and genes such as *IL1B[30], CXCL8[31], IFNG[31]*, and *TNF [32]* outlines a dense functional architecture, indicating that tumor-associated inflammation may be an orchestrated rather than a random process[33–35]. We propose that these axes represent critical maintenance hubs, where microbiota and immunity cooperate to perpetuate chronic inflammation.

Conversely, in PTT, functional organization was more diffuse. The association of *Helicobacter*, resting dendritic cells (cibersort_Dendritic cells resting), and regulatory genes such as *TGFB1* and *IL10* suggests an immunomodulated microenvironment capable of containing inflammation at subclinical levels[36]. The preservation of functional compartmentalization in PTT contrasts with the collapse observed in GAC, suggesting that tumor progression may involve the gradual dissolution of these regulatory barriers[37].

These observations are reinforced by topological analyses showing spatial overlap of activated B cells, immunosuppressive macrophages, inflammatory myeloid cells, and oral bacteria in GAC, versus organized segregation between adaptive responses and commensal microbiota in PTT[38, 39]. This structural opposition suggests that gastric cancer progression may be driven not only by inflammatory expansion but also by the loss of functional compartmentalization among immunity, inflammation, and microbial stimuli.

These findings have relevant clinical implications. The identification of microbiome-immune consortia organized around inflammatory genes in GAC points to potential therapeutic strategies targeting the disruption of these networks - for instance, through microbiota modulation or stromal reprogramming - aiming to restore local immune surveillance. Future capabilities to map inflammatory hotspots in the gastric microenvironment may guide more precise local or systemic therapies.

However, certain limitations must be acknowledged. Bulk RNA-based approaches do not permit single-cell spatial resolution, and inferences drawn from deconvolution and correlation analyses, while robust, require additional experimental validation. Moreover, the lack of longitudinal data limits the evaluation of the temporal dynamics of the observed networks.

Despite these limitations, this study provides a new perspective on the functional interaction among the microbiome, immunity, and gastric cancer progression. By demonstrating that gastric carcinogenesis is associated not merely with compositional changes but with the active formation of organized inflammatory networks, our findings propose new paradigms for the understanding and therapeutic targeting of the disease.

## Acknowledgments

The authors express their gratitude to CAPES (Coordenação de Aperfeiçoamento de Pessoal de Nível Superior) for providing a doctoral fellowship to R.M. da S. Mourão. We are also grateful to the High-Performance Computing Center (CCAD) at the Federal University of Pará for their support in computational resources. Furthermore, we acknowledge the Fundação Amazônia de Amparo a Estudos e Pesquisas (Fapespa) for the financial support that made this research possible.

## Data Availability

The datasets generated and analyzed during the current study are available from the corresponding author on reasonable request.

## Appendix I

**Figure 4.** Relative proportion of families and genera. (A) Relative abundance of predominant bacterial families; (B) Relative abundance of predominant bacterial genera. Each color represents a families or genera. Legends indicate the proportion of families or genera in GAC and PTT and Total

**Figure 5.** Relative proportion of cell. Composition of immune cell fractions across individual samples. Each color represents a cell type identified by deconvolution analysis. Legends indicate the proportion of the cell types in GAC and PTT and Total;

## References

1. Sathe, A., Grimes, S.M., Lau, B.T., Chen, J., Suarez, C., Huang, R.J., Poultsides, G., Ji, H.P.: Single-Cell Genomic Characterization Reveals the Cellular Reprogramming of the Gastric Tumor Microenvironment. Clinical Cancer Research. 26, 2640–2653 (2020). 10.1158/1078-0432.CCR-19-3231.

2. Jiang, X., Peng, L., Zhang, L., Li, Z., Zhou, T., Zhang, J., Li, W., You, W., Zhang, Y., Pan, K.: Gastric microbiota and its role in gastric carcinogenesis. Malignancy Spectrum. 1, 2–14 (2024). 10.1002/msp2.15.

3. Soneson, C., Love, M.I., Robinson, M.D.: Differential analyses for RNA-seq: transcript-level estimates improve gene-level inferences. F1000Res. 4, 1521 (2016). 10.12688/f1000research.7563.2.

4. Yong, X., Tang, B., Li, B.-S., Xie, R., Hu, C.-J., Luo, G., Qin, Y., Dong, H., Yang, S.-M.: Helicobacter pylori virulence factor CagA promotes tumorigenesis of gastric cancer via multiple signaling pathways. Cell Communication and Signaling. 13, 30 (2015). 10.1186/s12964-015-0111-0.

5. Mager, L.F., Krause, T., McCoy, K.D.: Interaction of microbiota, mucosal malignancies, and immunotherapy—Mechanistic insights. Mucosal Immunol. 17, 402–415 (2024). 10.1016/j.mucimm.2024.03.007.

6. Motamed, R., Jabbari, K., Sheikhbahaei, M., Ghazimoradi, M.H., Ghodsi, S., Jahangir, M., Habibi, N., Babashah, S.: Mesenchymal stem cells modulate breast cancer progression through their secretome by downregulating ten-eleven translocation 1. Sci Rep. 15, 6593 (2025). 10.1038/s41598-025-91314-3.

7. Belkaid, Y., Harrison, O.J.: Homeostatic Immunity and the Microbiota. Immunity. 46, 562–576 (2017). 10.1016/j.immuni.2017.04.008.

8. Ding, J.-T., Yang, K.-P., Zhou, H.-N., Huang, Y.-F., Li, H., Zong, Z.: Landscapes and mechanisms of CD8+ T cell exhaustion in gastrointestinal cancer. Front Immunol. 14, (2023). 10.3389/fimmu.2023.1149622.

9. Zeng, R., Gou, H., Lau, H.C.H., Yu, J.: Stomach microbiota in gastric cancer development and clinical implications. Gut. 73, 2062–2073 (2024). 10.1136/gutjnl-2024-332815.

10. Liu, Z., Zhang, D., Chen, S.: Unveiling the gastric microbiota: implications for gastric carcinogenesis, immune responses, and clinical prospects. Journal of Experimental & Clinical Cancer Research. 43, 118 (2024). 10.1186/s13046-024-03034-7.

11. Niikura, R., Hayakawa, Y., Nagata, N., Miyoshi-Akiayama, T., Miyabayashi, K., Tsuboi, M., Suzuki, N., Hata, M., Arai, J., Kurokawa, K., Abe, S., Uekura, C., Miyoshi, K., Ihara, S., Hirata, Y., Yamada, A., Fujiwara, H., Ushiku, T., Woods, S.L., Worthley, D.L., Hatakeyama, M., Han, Y.W., Wang, T.C., Kawai, T., Fujishiro, M.: Non-Helicobacter pylori Gastric Microbiome Modulates Prooncogenic Responses and Is Associated With Gastric Cancer Risk. Gastro Hep Advances. 2, 684–700 (2023). 10.1016/j.gastha.2023.03.010.

12. Noto, J.M., Peek, R.M.: The gastric microbiome, its interaction with Helicobacter pylori, and its potential role in the progression to stomach cancer. PLoS Pathog. 13, e1006573 (2017). 10.1371/journal.ppat.1006573.

13. Zavros, Y., Merchant, J.L.: The immune microenvironment in gastric adenocarcinoma. Nat Rev Gastroenterol Hepatol. 19, 451–467 (2022). 10.1038/s41575-022-00591-0.

14. Patro, R., Duggal, G., Love, M.I., Irizarry, R.A., Kingsford, C.: Salmon provides fast and bias-aware quantification of transcript expression. Nat Methods. 14, 417–419 (2017). 10.1038/nmeth.4197.

15. Love, M.I., Huber, W., Anders, S.: Moderated estimation of fold change and dispersion for RNA-seq data with DESeq2. Genome Biol. 15, 550 (2014). 10.1186/s13059-014-0550-8.

16. Wood, D.E., Lu, J., Langmead, B.: Improved metagenomic analysis with Kraken 2. Genome Biol. 20, 257 (2019). 10.1186/s13059-019-1891-0.

17. Barra, W.F., Sarquis, D.P., Khayat, A.S., Khayat, B.C.M., Demachki, S., Anaissi, A.K.M., Ishak, G., Santos, N.P.C., dos Santos, S.E.B., Burbano, R.R., Moreira, F.C., de Assumpção, P.P.: Gastric Cancer Microbiome. Pathobiology. 88, 156–169 (2021). 10.1159/000512833.

18. Newman, A.M., Liu, C.L., Green, M.R., Gentles, A.J., Feng, W., Xu, Y., Hoang, C.D., Diehn, M., Alizadeh, A.A.: Robust enumeration of cell subsets from tissue expression profiles. Nat Methods. 12, 453–457 (2015). 10.1038/nmeth.3337.

19. Finotello, F., Mayer, C., Plattner, C., Laschober, G., Rieder, D., Hackl, H., Krogsdam, A., Loncova, Z., Posch, W., Wilflingseder, D., Sopper, S., Ijsselsteijn, M., Brouwer, T.P., Johnson, D., Xu, Y., Wang, Y., Sanders, M.E., Estrada, M. V., Ericsson-Gonzalez, P., Charoentong, P., Balko, J., de Miranda, N.F. da C.C., Trajanoski, Z.: Molecular and pharmacological modulators of the tumor immune contexture revealed by deconvolution of RNA-seq data. Genome Med. 11, 34 (2019). 10.1186/s13073-019-0638-6.

20. Racle, J., Gfeller, D.: EPIC: A Tool to Estimate the Proportions of Different Cell Types from Bulk Gene Expression Data. Presented at the (2020). 10.1007/978-1-0716-0327-7_17.

21. Stewart, O.A., Wu, F., Chen, Y.: The role of gastric microbiota in gastric cancer. Gut Microbes. 11, 1220–1230 (2020). 10.1080/19490976.2020.1762520.

22. Battaglia, T.W., Mimpen, I.L., Traets, J.J.H., van Hoeck, A., Zeverijn, L.J., Geurts, B.S., de Wit, G.F., Noë, M., Hofland, I., Vos, J.L., Cornelissen, S., Alkemade, M., Broeks, A., Zuur, C.L., Cuppen, E., Wessels, L., van de Haar, J., Voest, E.: A pan-cancer analysis of the microbiome in metastatic cancer. Cell. 187, 2324–2335.e19 (2024). 10.1016/j.cell.2024.03.021.

23. Engstrand, L., Graham, D.Y.: Microbiome and Gastric Cancer. Dig Dis Sci. 65, 865–873 (2020). 10.1007/s10620-020-06101-z.

24. Li, X., Pan, K., Vieth, M., Gerhard, M., Li, W., Mejías-Luque, R.: JAK-STAT1 Signaling Pathway Is an Early Response to Helicobacter pylori Infection and Contributes to Immune Escape and Gastric Carcinogenesis. Int J Mol Sci. 23, 4147 (2022). 10.3390/ijms23084147.

25. Zhang, X., Ren, B., Liu, B., Wang, R., Li, S., Zhao, Y., Zhou, W.: Single-cell RNA sequencing and spatial transcriptomics reveal the heterogeneity and intercellular communication of cancer-associated fibroblasts in gastric cancer. J Transl Med. 23, 344 (2025). 10.1186/s12967-025-06376-8.

26. Mak, T.K., Li, X., Huang, H., Wu, K., Huang, Z., He, Y., Zhang, C.: The cancer-associated fibroblast-related signature predicts prognosis and indicates immune microenvironment infiltration in gastric cancer. Front Immunol. 13, (2022). 10.3389/fimmu.2022.951214.

27. Zhao, H., Wu, L., Yan, G., Chen, Y., Zhou, M., Wu, Y., Li, Y.: Inflammation and tumor progression: signaling pathways and targeted intervention. Signal Transduct Target Ther. 6, 263 (2021). 10.1038/s41392-021-00658-5.

28. Batlle, E., Massagué, J.: Transforming Growth Factor-β Signaling in Immunity and Cancer. Immunity. 50, 924–940 (2019). 10.1016/j.immuni.2019.03.024.

29. Park, C.H., Hong, C., Lee, A., Sung, J., Hwang, T.H.: Multi-omics reveals microbiome, host gene expression, and immune landscape in gastric carcinogenesis. iScience. 25, 103956 (2022). 10.1016/j.isci.2022.103956.

30. Niikura, R., Hayakawa, Y., Nagata, N., Miyoshi-Akiayama, T., Miyabayashi, K., Tsuboi, M., Suzuki, N., Hata, M., Arai, J., Kurokawa, K., Abe, S., Uekura, C., Miyoshi, K., Ihara, S., Hirata, Y., Yamada, A., Fujiwara, H., Ushiku, T., Woods, S.L., Worthley, D.L., Hatakeyama, M., Han, Y.W., Wang, T.C., Kawai, T., Fujishiro, M.: Non-Helicobacter pylori Gastric Microbiome Modulates Prooncogenic Responses and Is Associated With Gastric Cancer Risk. Gastro Hep Advances. 2, 684–700 (2023). 10.1016/j.gastha.2023.03.010.

31. Duizer, C., Salomons, M., van Gogh, M., Gräve, S., Schaafsma, F.A., Stok, M.J., Sijbranda, M., Kumarasamy Sivasamy, R., Willems, R.J.L., de Zoete, M.R.: Fusobacterium nucleatum upregulates the immune inhibitory receptor PD-L1 in colorectal cancer cells via the activation of ALPK1. Gut Microbes. 17, (2025). 10.1080/19490976.2025.2458203.

32. Dharmani, P., Strauss, J., Ambrose, C., Allen-Vercoe, E., Chadee, K.: Fusobacterium nucleatum Infection of Colonic Cells Stimulates MUC2 Mucin and Tumor Necrosis Factor Alpha. Infect Immun. 79, 2597–2607 (2011). 10.1128/IAI.05118-11.

33. Gobert, A.P., Wilson, K.T.: Induction and Regulation of the Innate Immune Response in Helicobacter pylori Infection. Cell Mol Gastroenterol Hepatol. 13, 1347–1363 (2022). 10.1016/j.jcmgh.2022.01.022.

34. Zheng, W., Wang, Y., Sun, H., Bao, S., Ge, S., Quan, C.: The role of Fusobacterium nucleatum in macrophage M2 polarization and NF-κB pathway activation in colorectal cancer. Front Immunol. 16, (2025). 10.3389/fimmu.2025.1549564.

35. Zhang, J., Hu, C., Zhang, R., Xu, J., Zhang, Y., Yuan, L., Zhang, S., Pan, S., Cao, M., Qin, J., Cheng, X., Xu, Z.: The role of macrophages in gastric cancer. Front Immunol. 14, (2023). 10.3389/fimmu.2023.1282176.

36. Kao, J.Y., Zhang, M., Miller, M.J., Mills, J.C., Wang, B., Liu, M., Eaton, K.A., Zou, W., Berndt, B.E., Cole, T.S., Takeuchi, T., Owyang, S.Y., Luther, J.: Helicobacter pylori Immune Escape Is Mediated by Dendritic Cell-Induced Treg Skewing and Th17 Suppression in Mice. Gastroenterology. 138, 1046–1054 (2010). 10.1053/j.gastro.2009.11.043.

37. Boesch, M., Horvath, L., Baty, F., Pircher, A., Wolf, D., Spahn, S., Straussman, R., Tilg, H., Brutsche, M.H.: Compartmentalization of the host microbiome: how tumor microbiota shapes checkpoint immunotherapy outcome and offers therapeutic prospects. J Immunother Cancer. 10, e005401 (2022). 10.1136/jitc-2022-005401.

38. Lee, S.H., Lee, D., Choi, J., Oh, H.J., Ham, I.-H., Ryu, D., Lee, S.-Y., Han, D.-J., Kim, S., Moon, Y., Song, I.-H., Song, K.Y., Lee, H., Lee, S., Hur, H., Kim, T.-M.: Spatial dissection of tumour microenvironments in gastric cancers reveals the immunosuppressive crosstalk between CCL2+ fibroblasts and STAT3-activated macrophages. Gut. 74, 714–727 (2025). 10.1136/gutjnl-2024-332901.

39. Wang, R., Song, S., Qin, J., Yoshimura, K., Peng, F., Chu, Y., Li, Y., Fan, Y., Jin, J., Dang, M., Dai, E., Pei, G., Han, G., Hao, D., Li, Y., Chatterjee, D., Harada, K., Pizzi, M.P., Scott, A.W., Tatlonghari, G., Yan, X., Xu, Z., Hu, C., Mo, S., Shanbhag, N., Lu, Y., Sewastjanow-Silva, M., Fouad Abdelhakeem, A.A., Peng, G., Hanash, S.M., Calin, G.A., Yee, C., Mazur, P., Marsden, A.N., Futreal, A., Wang, Z., Cheng, X., Ajani, J.A., Wang, L.: Evolution of immune and stromal cell states and ecotypes during gastric adenocarcinoma progression. Cancer Cell. 41, 1407–1426.e9 (2023). 10.1016/j.ccell.2023.06.005.

