## Supplementary figures and images for "Immune Remodeling and Dysbiosis May Distinguish the Microenvironments of Gastric Adenocarcinoma and Peritumoral Tissue"

### figure 4.tiff

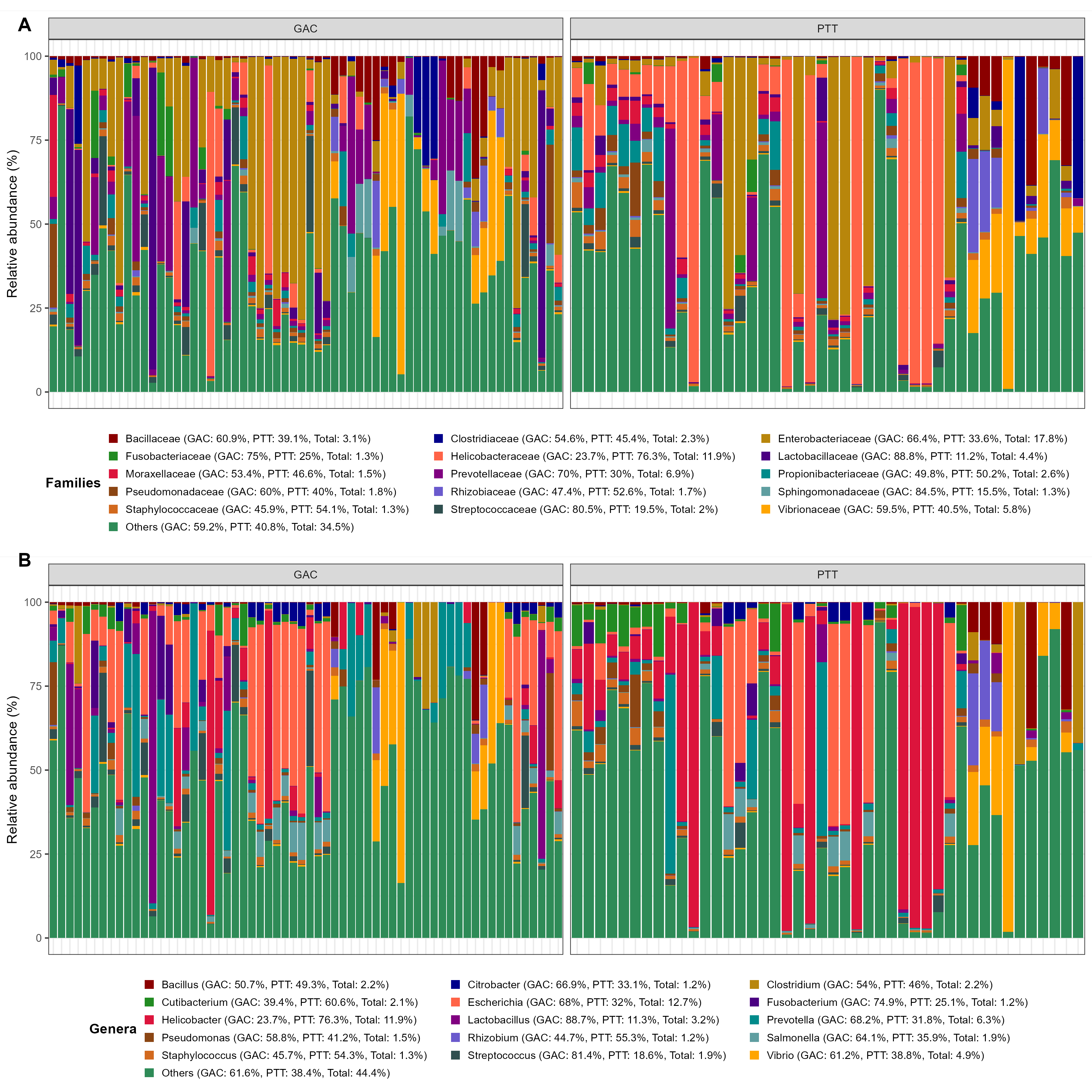

### figure 5.tiff

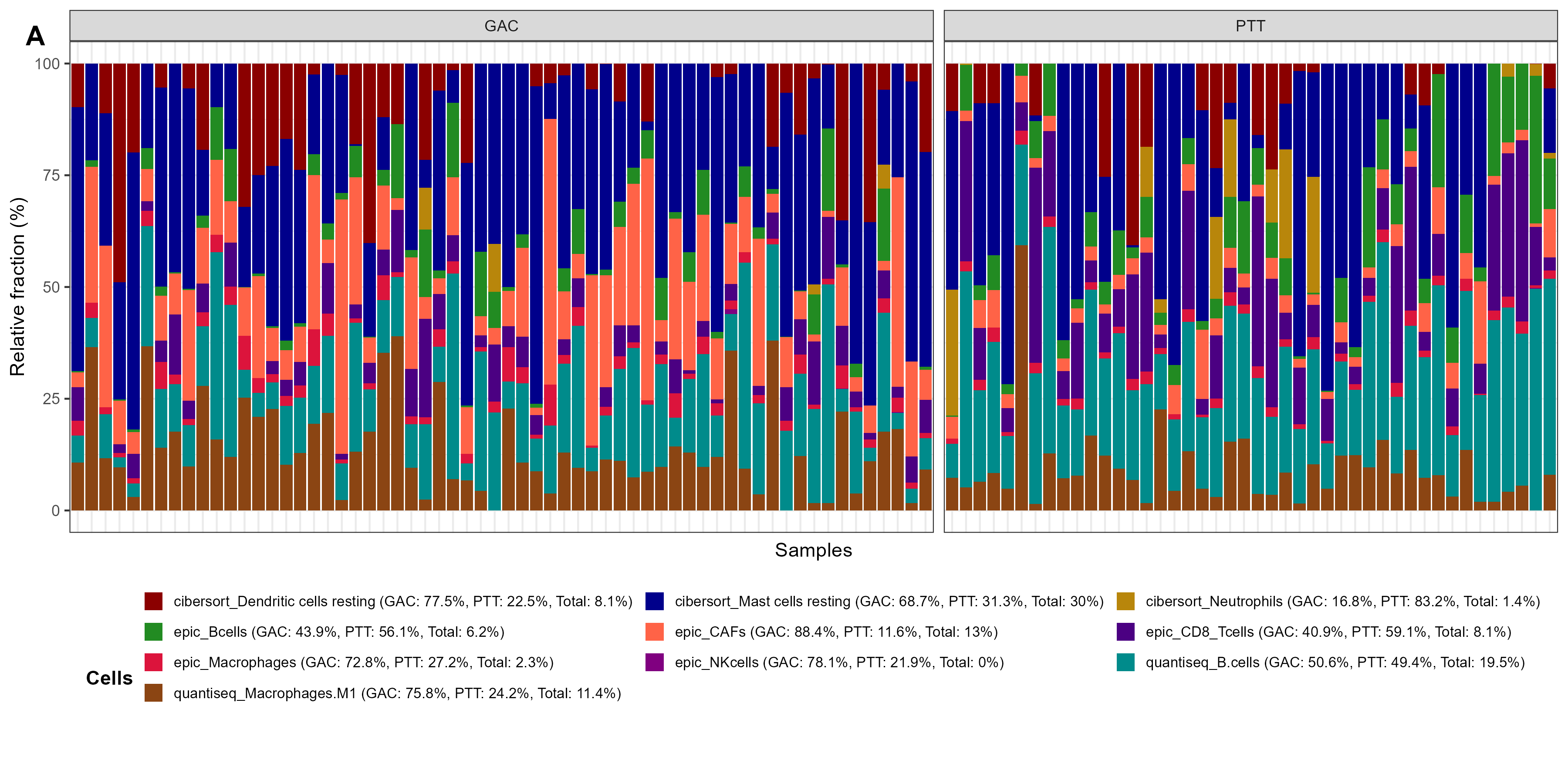
